# Rapid and accurate taxonomic classification of cpn60 amplicon sequence variants

**DOI:** 10.1101/2023.01.29.526108

**Authors:** Qingyi Ren, Janet E. Hill

**Affiliations:** Department of Veterinary Microbiology, University of Saskatchewan

## Abstract

The “universal target” region of the gene encoding the 60 kDa chaperonin protein (cpn60, also known as groEL or hsp60) is a proven sequence barcode for bacteria and a useful target for marker gene amplicon-based studies of complex microbial communities. To date, identification of cpn60 sequence variants from microbiome studies has been accomplished by alignment of queries to a reference database. Naïve Bayesian classifiers including the RDP classifier offer an alternative identification method that provides variable rank classification and shorter analysis times. We curated a set of cpn60 barcode sequences to train the RDP classifier and tested its performance on data from previous human microbiome studies. Results showed that sequences accounting for 79%, 86% and 92% of the observations (read counts) in saliva, vagina and infant stool microbiome data sets were classified to the species rank. We also established a threshold confidence value of 0.6 at phylum rank for filtering non-target amplicon sequences from study data and demonstrated that trimming the training sequences to match the lengths of the queries is not needed for accurate cpn60 sequence classification. Successful implementation of a naïve Bayesian classifier for cpn60 sequences will facilitate future microbiome studies and open opportunities to integrate cpn60 amplicon sequence identification into existing analysis pipelines.

## INTRODUCTION

High throughput, massively parallel sequencing of PCR amplified marker gene sequences (barcodes) remains an important technique for determining the taxonomic composition of complex microbial communities. While variable regions of the 16S rRNA gene remain the most widely targeted markers for bacteria, high resolution microbiome profiling has also been achieved using partial cpn60 sequences. The cpn60 “universal target” region (corresponding to nucleotides 274-828 of the *E. coli* cpn60 gene, also known as hsp60 or groEL) has been demonstrated to be a preferred barcode for bacteria [1] and due to its conservation in bacteria and eukaryotes (and some archaea), it can provide simultaneous detection of bacteria and fungi in microbial communities [2]. cpn60 barcode sequences are excellent predictors of whole genome sequence relationships [3–5] and can be used for sub-species resolution of bacterial taxa [6–9]. The availability of “universal” PCR primers [10,11] targeting the cpn60 barcode and a manually curated database of reference sequences (cpnDB [12]) have made this sequence an attractive target for diagnostics [13–15], characterization of new species [16–18] and taxonomic profiling of a variety of microbial communities, including microbiota of human and animal body sites [19–24], and environmental samples [25–27].

Regardless of the marker gene targeted in an amplicon sequencing study, taxonomic identification of the resulting sequence requires a reference database appropriate for the environment under investigation and a robust method for sequence comparison that results in accurate identification. A wide range of alignment methods have been developed for comparing microbiome derived sequences to reference databases. For alignment of cpn60 barcode sequences amplified from microbiomes, wateredBLAST [28] was developed. wateredBLAST uses a combination of BLASTn and Smith-Waterman alignments and results in the identification of the “nearest neighbour” in the reference database for each query based on percent sequence identity over the entire length of the query. A limitation of this approach, and any other alignment-based method, is that only species level identification is reported even if the percent identity is low; two amplicon sequences with different levels of identity to the same nearest neighbour end up with the same identification. This is a larger issue for alignment of data from environments whose constituents are not well represented in the reference database. Another disadvantage of alignment methods in general is that they can be slow and expensive in terms of computational resources, requiring large amounts of memory. This is increasingly problematic as microbiome studies expand in size and scope, with a corresponding increase in the amount of sequence data produced.

Sequence classifiers offer an alternative to alignment methods and have two potential advantages: they tend to be faster since they reduce queries and databases to unique “words” (kmers), and they allow variable rank classifications since taxonomic lineage information is included in the reference data. Users can decide which ranks to accept based on the confidence value associated with each result. Several different classifiers have been developed with the naive Bayesian classifier, RDP Classifier [29], being perhaps the most widely used. The RDP Classifier was originally designed for 16S rRNA data from prokaryotes, but if training data is available, it can be used for other targets, such as large-subunit RNA (LSU) in fungi [30] cytochrome oxidase subunit 1 (CO1) in animals [31], and rbcL and ITS2 sequences in plants [32]. The success of sequence classification depends upon the availability of high-quality reference data with complete taxonomic lineage information. Incorrectly annotated reference sequences can lead to false positive classification results, while gaps in coverage of the reference database relative to the environment under study can lead to failure to classify beyond the highest taxonomic ranks.

The cpnDB reference database is manually curated and focuses on incorporation of bacterial type strain data to ensure robust landmarks for sequence identification [12]. The establishment of amplicon sequence variant (ASV) calling for cpn60 barcode sequences and the demonstration that as little as 150 bp from the 5’ end of the barcode region can provide species level identification [33] have resulted in improvements to analysis of cpn60 sequence data from microbiome studies, however, reliance on alignment methods for taxonomic identification remains a limitation. The objectives of the current study were to evaluate use of the RDP classifier for taxonomic assignment of cpn60 ASV data and to provide a trained classifier to the research community. We evaluated performance of the classifier in terms of accuracy and processing time and compared its results to those from wateredBLAST alignment using data from previously conducted studies of human salivary, vaginal, and infant stool microbiomes.

## MATERIALS AND METHODS

### Generation of training files

The RDP classifier (version 2.13) provided within RDPTools was used in this study. All group I reference nucleotide universal target (UT) sequences in the cpnDB reference database (accessed May 2022) were used to generate the training files. From the reference database, the name of each taxon (genus+species) and cpn60 UT sequence (fasta files) were obtained. The full taxonomic lineage for each taxon was obtained from the NCBI taxonomy database using the *taxarank* function in taxonomy_ranks [34,35]. For each taxon, any taxonomic rank without a corresponding entry returned a result of “NA”, which was replaced with “-” to make the resulting taxonomy table interpretable in subsequent steps. For any taxon that returned more than one *taxarank* result (e.g. where a eukaryote and a prokaryote share the same genus+species name), the eukaryotic lineage was then manually deleted from the taxonomy table. Similarly, taxa without a lineage that extended to the species level were removed from the taxonomy table and from the sequence data file. Python scripts (*Lineage2taxTrain* and *addFullLineage*) for creating “ready to train” taxonomy table and fasta files were obtained from https://github.com/GLBRC-TeamMicrobiome/python_scripts. The *Lineage2taxTrain* python script was then applied to the modified taxonomy table to produce the “ready to train” taxonomy file. For the fasta files, the *rm-dupseq* function of the classifier was used to remove any duplicate sequences (since these can inflate results during classification performance testing), then the *addFullLineage* script was used to insert lineage information into the definition line of each sequence. Finally, *Classifier*.*jar* was used to generate the training files from the two ready to train files, and the rRNAClassifier.properties file was downloaded from https://github.com/rdpstaff/classifier/blob/master/samplefiles/rRNAClassifier.properties and added in the output directory.

### Testing and evaluating the performance of the classifier

Initial testing of the trained classifier was performed by using the training sequences as queries and evaluating the results in terms of whether sequences were correctly classified as themselves.

Leave-one-out-testing (LOOT) was performed to evaluate the accuracy of the classifier using the built-in leave-one-out command within the RDP classifier with the LOOT default confidence cutoff of 0.9. Classifiers trained on different 5’-anchored lengths of cpn60 barcode sequence (50 bp, 150 bp, 300 bp, 450 bp and full length) were used in LOOT, and the accuracy of results for each query length were compared.

### Classification of microbiome sequence data and comparison with wateredBLAST

ASV sequences (150 bp) from three previously collected microbiome data sets (saliva (6737 ASV from 89 samples)[36], vagina (1950 ASV from 283 samples)[37], and infant stool (4153 ASV from 584 samples)[38]) were analyzed using the trained classifier with the default confidence value of ≥0.8 to assign a rank. To evaluate the accuracy of the classifier results, the same queries were identified using wateredBLAST for comparison. The same set of cpn60 sequences used to train the classifier was used as the reference database for wateredBLAST. Results from wateredBLAST and the classifier were compared based on species level identification/classification results and time taken for the analysis. Species rank confidence and wateredBLAST percent identity results for each sample type were visualized XY density plots using geom_density_2d and ggplot2 in R (version 4.2.1).

To test whether having the query the same length as the sequences in the training set influenced the classification results, the classifier was trained on the reference data set trimmed to the first 150 bp. The stool microbiome ASV sequences (150 bp) were then analyzed using either the Classifier trained on 150 bp or full-length (∼555 bp) cpn60 barcode sequences and results were compared using Spearman’s correlation test in GraphPad Prism v 9.5.0.

All classifier results were parsed with a custom python script to extract confidence values for each taxonomic rank for analysis.

### Establishing classifier thresholds for identification of non-cpn60 sequences

To determine a recommended cutoff (taxonomic rank and confidence level) for the classifier to filter out non-cpn60 sequences, two values were chosen for further assessment based on visual inspection of initial results: 0.6 and 0.8 at the phylum level. In order to choose which one was a better indicator of non-cpn60 sequences, specificity and sensitivity were calculated using wateredBLAST as the gold standard since it has been demonstrated that the minimum identity of cpn60 sequences to any cpnDB reference sequence is 55% [39]. Sensitivity was calculated as: true positive/(true positive + false negative) and specificity was calculated as: true negative/(true negative + false positive).

## RESULTS AND DISCUSSION

### Training the classifier

The cpnDB reference database of 17,815 sequences was selected as the initial training data. After removing duplicates and errors, 17,778 entries were included. The *taxarank* script was then applied to obtain the full taxonomic lineage of all taxa, which generated 17,793 entries in the resulting taxonomy table because some taxa yielded multiple entries. Duplicate entries were removed manually, leaving 17,778 entries. Taxa that were not labeled to the species level (e.g., *Staphylococcus* sp.) were removed leaving 16,416 entries. The *rm-dupseq* function of the RDP Classifier was used to remove duplicate sequences in the training set, resulting in 11,001 unique sequences to train the classifier. This is obviously much smaller than commonly used 16S rRNA training sets, such as the Silva database, which contains 128,884 bacterial, 2,846 archaeal, and 14,871 eukaryotic sequences as of release 138.1 [40]. Even though cpn60 training set has fewer sequences, it has substantial taxonomic breadth as a result of the cpnDB curation strategy, which emphasizes broad coverage of taxa rather than many redundant entries for individual species [12].

### Initial testing of the classifier

After training, the functionality of the classifier and the validity of the training files were tested by using the training set as queries. The training set contained 11,001 sequences, and 9,849 (89.5%) of these were classified to the species rank with a confidence level of 1.0 (i.e., they were classified as themselves as expected). Out of the remaining 1152 sequences, 799 had a confidence value of 0.8-0.99 at the species rank. Out of the 353 sequences which were not classified confidently at the species rank, 321 (90.9%) were classified at the genus rank.

### Leave-one-out testing

In leave-one-out testing (LOOT), each sequence is removed in turn from the training set and then classified based on the remaining sequences. To determine the effect of sequence length on classification accuracy, LOOT was performed with 5’ anchored sequence lengths of 50, 150, 300, 450 bp and full-length (∼555 bp) cpn60 barcode (Figure 1). Regardless of length, >90% of queries were accurately classified to the ranks of kingdom, phylum and class. At lower taxonomic ranks, the 50 bp sequences were classified markedly more poorly than longer sequences.

**Figure 1.**
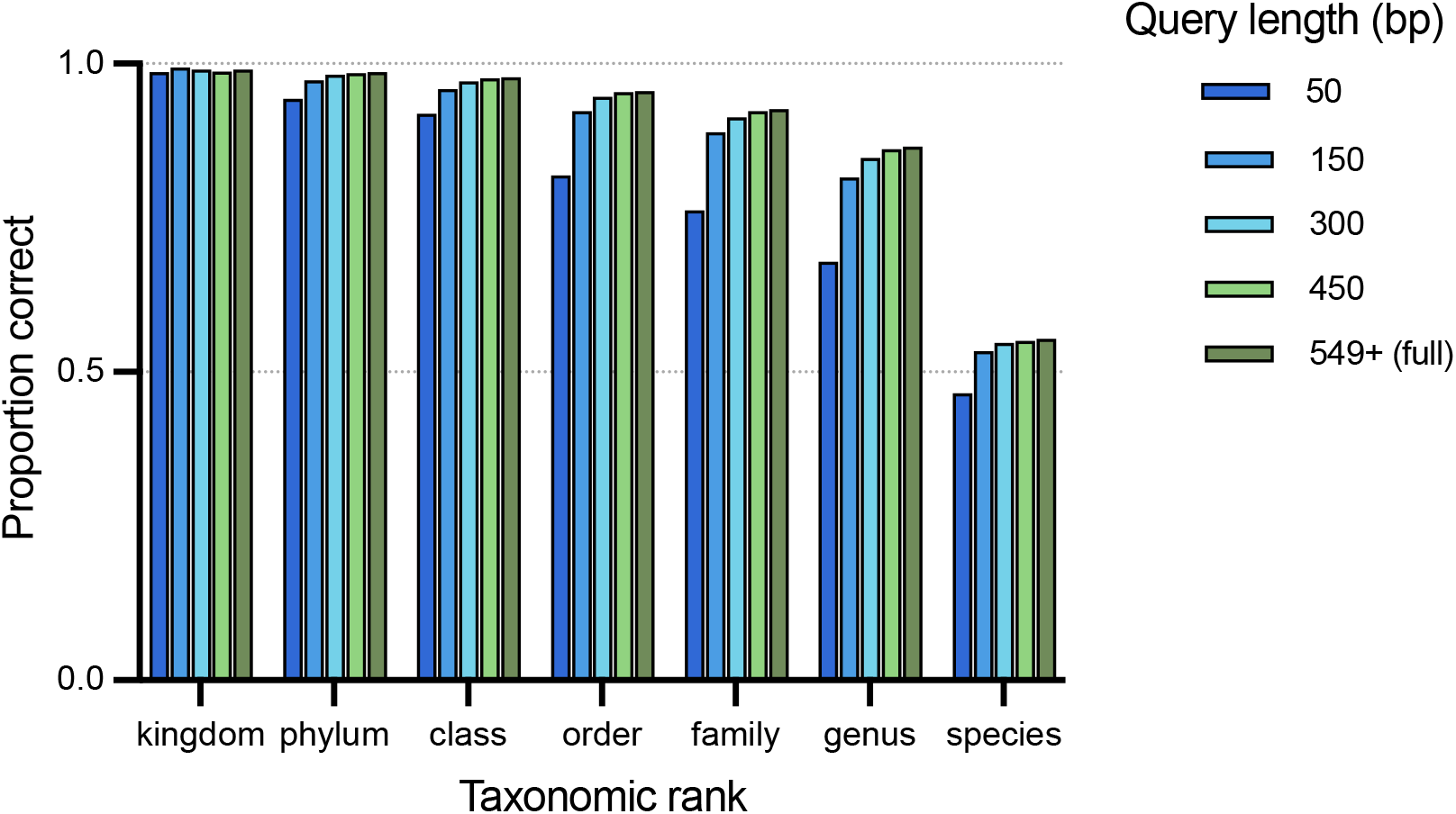
Effects of sequences lengths on LOOT results. Five different 5’ anchored lengths of the same set of 11,001 sequences were assessed.

An approximately 30% of drop in the proportion of accurately classified sequences was observed at the species rank relative to genus rank. Any species that was unique in the training set could obviously not be correctly classified in LOOT. Accurate species level classification was only possible in cases where there were multiple strains of individual species with non-identical cpn60 sequences in the training set. In some cases, the complete database did contain multiple records for a species but if the cpn60 sequences were identical they were removed by the remove-duplicate step in preparing the training files. If there were multiple entries for a species with non-identical cpn60 sequences in the training set, accurate species level classification was possible, which was the case for just over half of the queries in the LOOT experiment (Figure 1). Sequence lengths of ≥150 bp all provided accurate results at the genus rank for at least 80% of queries, and the proportion of accurately classified sequences increased as the query length increased (Figure 1). As expected, the proportion of accurately classified sequences also increased at higher taxonomic ranks. Improved accuracy with longer query lengths was expected given the nature of a naive-Bayesian classifier, where longer sequences generate more “words” for the classifier to look up. However, the improvement in accuracy for lengths beyond 150 bp was minimal. This observation is consistent with previous work showing that using alignment methods, 150 bp from the 5’ end of the barcode region is sufficient for unambiguous identification [41].

### Classification of human microbiome sequence data

To compare the classifier and wateredBLAST for species level identification of cpn60 ASV from microbiomes, we analyzed ASV sequences from previously conducted studies focused on salivary, vaginal and infant stool microbiomes. ASV sequences were aligned to the training sequences with wateredBLAST or identified using the classifier with results plotted as percent identity to a reference sequence or confidence at species rank, respectively (Figure 2). ASV identification was more rapid with the classifier than wateredBLAST for each set of queries: stool (28.1 s *vs*. 86.4 s), vagina (49.3 s *vs*. 198.1 s) and infant stool (72.8 s *vs*. 323.6 s).

**Figure 2.**
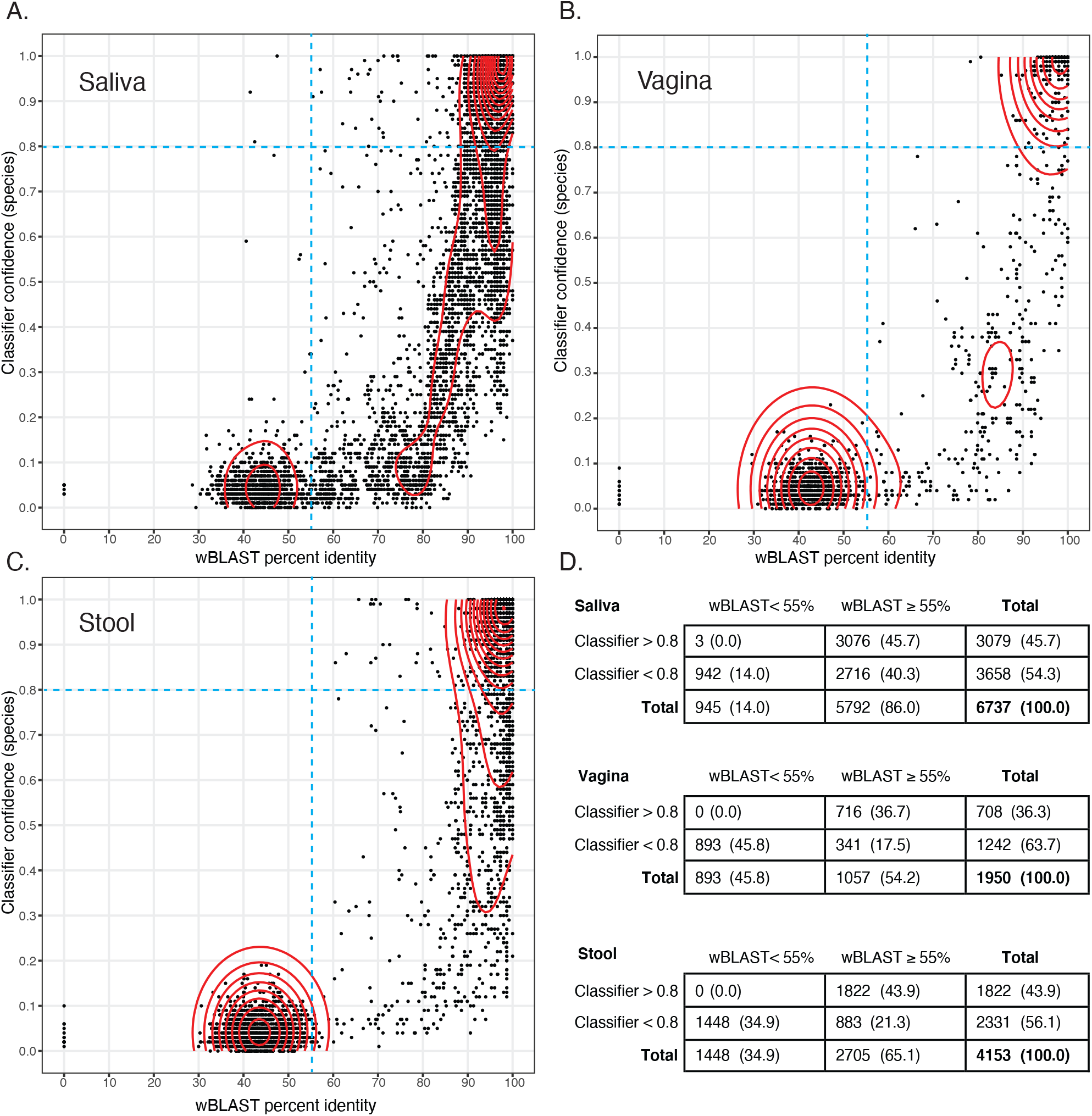
Taxonomic identification of 150 bp cpn60 ASV from saliva (A), vagina (B) and infant stool (C) microbiomes by wateredBLAST (abscissa, % identity) and the classifier (ordinate, confidence at species rank). Density of data points is indicated by red contour lines. Broken blue lines indicate two thresholds: ≥55% sequence identity for wateredBLAST (the established cut-off for non-cpn60 sequences) and 0.8 for the classifier (the default confidence threshold). Number (percent) of ASV in each quadrant of the plots are shown in (D).

We divided the x-y plots into four quadrants based on two values: ≥55% for wateredBLAST and ≥0.8 for the classifier (Figure 2). The left two quadrants consist of sequences that have a sequence identity of <55%, which according to our past experience, are not cpn60 [39]. As expected, queries with wateredBLAST scores of <55% had correspondingly low classifier confidence values at the species rank, generally <0.2 (Figure 2). Interestingly there were three queries in the salivary microbiome data that gave wateredBLAST scores <55% but were classified confidently (>0.8) at the species level (Figure 2A). Despite the low wateredBLAST identities, the two methods agreed on the identification of these three ASV as *Rothia mucilaginosa, Selenomoas infelix* and *Centipeda periodontii*.

The right top quadrant in each plot in Figure 2 includes ASV that are cpn60 based on the ≥55% wateredBLAST cutoff criterion and are confidently classified at species rank by the classifier. For all microbiome data sets this quadrant accounts for the majority ASV that met the wateredBLAST criterion to be identified as a cpn60 sequence. Additionally, the majority of these ASV were identified as the same species by wateredBLAST and the classifier. Among ASV classified with confidence ≥0.8 at the species rank and ≥90% identity by wateredBLAST, 2735/2931 (93.3%) of saliva ASV, 683/705 (96.9%) of vaginal ASV, and 1617/1780 (90.8%) of stool ASV had the same species identification result. In cases where the wateredBLAST and classifier species results did not agree there was agreement on genus, although in a handful of cases there disagreement was related to a recent name change (e.g., *Atlantibacter* vs. *Escherichia*).

The lower right quadrants of the density plots in Figure 2 contain ASV queries that are cpn60 (≥55% identical by wateredBLAST) but are not confidentially classified at the species rank by the classifier. However, these sequences were classified confidently at higher ranks. 1024/2716 (37.7%) of these lower right quadrant ASV sequences from the saliva microbiome, 56/341 (16.4%) ASV sequences from the vaginal microbiome, and 435/883 (49.3%) ASV sequences from the infant stool microbiome were classified confidently at the genus rank and the remainder were classified at higher ranks.

To identify factors that contribute to a high wateredBLAST percent identity and a relatively low species rank classifier confidence, we examined results for saliva microbiome query sequences that were: identified with a percent identity of ≥90% by wateredBLAST and classified with a confidence level of <0.8 at species rank by the classifier. These queries were further divided into two categories: those that were classified with a confidence level of ≥0.8 at genus rank by the classifier (query type A, 887 ASV), and those that were classified with a confidence level of <0.8 at genus level by the classifier (query type B, 163 ASV). The number of relevant sequences in the training set corresponding to each query might be expected to influence classifier confidence, i.e. having few representatives of a taxon may lead to low confidence and thus queries belonging to a poorly represented taxon are more likely to receive low confidence scores. Conversely, having many closely related sequences representing a taxon might also result in low confidence values because the classifier cannot distinguish between several likely matches. This latter scenario likely explains why some queries were not classified with confidence as themselves in our initial testing. To investigate both possible explanations for the different qualities of classification results for ASV that were ≥90% identical to something in the database by wateredBLAST but not classified at the species rank, we determined the number of records in cpnDB corresponding to the putative identities of the Type A and B queries (Table 1). Interestingly there was no significant difference between the average number of records per genus for Type A or B queries (Mann Whitney, P=0.0668), suggesting that the confidence of classification is not completely dependent on the number of representative in the database.

**Table 1.** Database coverage for saliva ASV queries that were ≥90% identical their nearest database neighbour by wateredBLAST, but classified at species level with confidence <0.8. Type A queries were classified with confidence >0.8 at the genus rank, whereas Type B queries were classified with confidence <0.8 at genus rank.

| Query type | No. of ASV | No. of genera | cpnDB records per genus |  |  |  |
| --- | --- | --- | --- | --- | --- | --- |
|  |  |  | Min | Max | Median | Average |
| Type A | 887 | 26 | 1 | 213 | 18.5 | 46.2 |
| Type B | 163 | 23 | 1 | 211 | 9 | 26.7 |

### Determining a classifier threshold to filter out non-cpn60 sequences

All “universal” primer PCR amplifications of marker genes will result in some level of non-target amplification, and the amount of those contaminating sequences generated will be influenced by factors including overall target abundance and the presence of host DNA in the case of host-associated microbiome samples [39]. How these off-target sequences are handled in the bioinformatic pipeline following microbiome amplicon library sequencing is variable, and many studies do not include a distinction between true target sequences that are “unidentified” and non-target sequences. Extensive analysis of cpn60 ASV sequences from previous microbiome studies and observations of cpn60 sequences from prokaryotes and eukaryotes suggest that the minimum similarity between any two known cpn60 sequences is ∼55% [12,39].

All the ASV with ≥55% identity to a sequence in cpnDB also had a phylum rank classification with >0.8 confidence, however, not all ASV with a phylum rank confidence <0.8 were <55% identical to a cpnDB sequence. This result demonstrates that confidence <0.8 at the phylum rank is not an ideal classifier-based criterion for identification of non-cpn60 sequences since it would filter out true cpn60 sequences. Visual inspection of classifier results from the microbiome ASV data suggested that a phylum rank confidence value of 0.6 was a reasonable alternative. To further evaluate these potential thresholds for identification of true cpn60 sequences, we performed a sensitivity and specificity analysis using percent sequence identity by wateredBLAST as the gold standard test. Sensitivity and specificity were calculated for both proposed threshold values for each of the human microbiome query sets (Table 2). Specificity was higher for phylum confidence of ≥0.8 than ≥0.6 in all cases, but sensitivity was higher for ≥0.6 than ≥0.8, indicating that using a confidence of ≥0.8 at the phylum rank would be more likely to result in screening out true cpn60 sequences. In microbiome studies aimed at cataloging diversity, sensitivity should be prioritized over specificity since this reduces the chances of novel cpn60 sequences being screened out (false negatives), therefore 0.6 at phylum rank is recommended for filtering cpn60 ASV data.

**Table 2.** Sensitivity and specificity of two potential classifier cutoffs for identification of true cpn60 sequences (defined as ≥55% identical to nearest database neighbour by wateredBLAST).

| Classifier criterion |  | Saliva | Stool | Vagina | Average |
| --- | --- | --- | --- | --- | --- |
| Phylum confidence $\geq 0.6$ | Sensitivity | 0.853 | 0.974 | 0.913 | 0.913 |
|  | Specificity | 0.977 | 0.990 | 0.998 | 0.988 |
| Phylum confidence $\geq 0.8$ | Sensitivity | 0.788 | 0.947 | 0.826 | 0.895 |
|  | Specificity | 0.993 | 1.00 | 0.999 | 0.996 |

Once a cut-off was established, the results from the classifier for all three human microbiome data sets were re-examined with non-cpn60 sequences filtered out (Figure 3). Filtration using the phylum confidence ≥0.6 criterion retained 4960/6737, 967/1950, 2648/4153 ASV sequences in the saliva, vagina and infant stool microbiome data sets, respectively. In all three environments, the majority of 150 bp ASV queries were classified confidently at species level: 62%, 74% and 69% of saliva, vagina and stool ASV, respectively (Figure 3). When the relative abundance of the ASV in the microbiome data sets was considered, species level classification was achieved for 79%, 86% and 92% of the ASV observations (read counts) in the saliva, vagina and infant stool microbiomes, indicating that the more poorly classified sequences were also much less abundant in these microbiomes (Figure 3). Results varied slightly among the three environments, with a larger proportion of the vagina and stool derived ASV sequences classified at the species rank compared to ASV sequences from the saliva microbiome, likely reflecting the representation of these environments in cpnDB and the extent to which organisms from these environments have been characterized and named. In fact, several of the most abundant but poorly classified ASV from the saliva microbiome were identical to unidentified cpn60 sequences from metagenomic studies of the oral microbiome.

**Figure 3.**
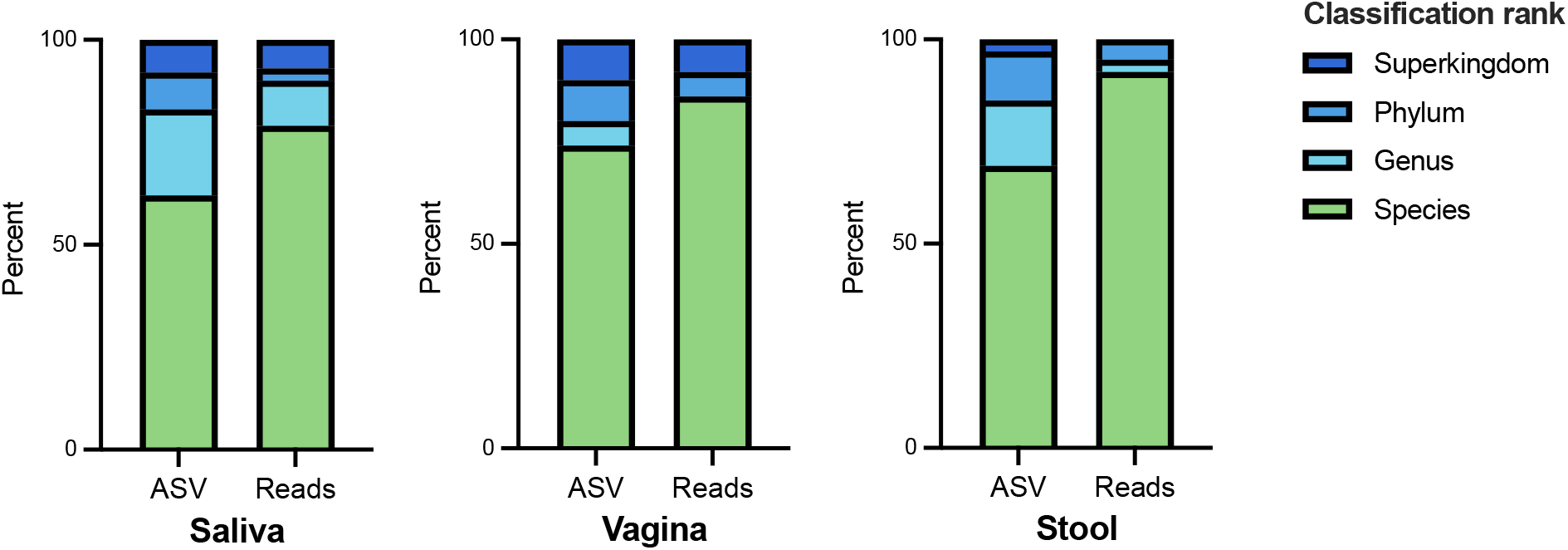
Percentage of cpn60 ASV sequences and proportion of the microbiome (percent of read counts across all samples) classified with a confidence level of >0.8 to the rank of species, genus, phylum and superkingdom. Results are shown for saliva (4960 ASV from 89 samples, 5.8M reads), vagina (967 ASV from 283 samples, 1.5M reads), and infant stool (2648 ASV from 584 samples, 2.4M reads) microbiomes.

### Effect of training sequence length on classifier results

The developers of the RDP classifier recommend using training sequences corresponding in length to the queries [42]. For example, if the data to be classified is V3-V4 of the 16S rRNA gene, better accuracy will be obtained if the database of full-length 16S rRNA gene sequences is trimmed to include only the V3-V4 region prior to training. We tested whether keeping the training set the same length as the queries would affect the results of the cpn60 classifier. We classified 150 bp stool microbiome ASV using the classifier trained on either 150 bp or full-length training sequences. Species rank confidence values were compared for each ASV and the results were highly correlated (Spearman’s correlation test, r = 0.9213, p <0.0001) suggesting no significant difference in performance when the training sequences were trimmed to the length of the queries. Typical 16S rRNA amplicon sequencing targets short variable regions that may account for a small proportion of the length of the 16S rRNA gene. For example, the V3 region is ∼175 bp of the ∼1.5 kb 16S rRNA gene [43], compared to the 150 bp cpn60 sequence that accounts for proportionately more of the ∼555 bp “universal target” region of cpn60. In addition, the presence of multiple conserved and variable regions within the 16S rRNA gene sequence (compared to the more uniform diversity across the cpn60 target [1]) may present a further obstacle for the classifier.

In conclusion, the RDP Classifier gives comparable results to wateredBLAST for identification of microbiome derived cpn60 sequences of at least 150 bp and provided rapid species rank classification for ASV accounting for up to 92% of three different human microbiome data sets. The proportion of ASV classified to species rank will no doubt be improved with continuing expansion of cpnDB, by strategic enrichment of reference data for microbiomes of interest, and continuing efforts to characterize and name novel bacteria from diverse environments.

## Acknowledgements

The authors are grateful to John Quensen for his informative on-line instructions on training the RDP Classifier. Thanks to Jay Rabari for troubleshooting python scripts. This project was funded by a Natural Sciences and Engineering Research Council of Canada (NSERC) Discovery Grant.

## Data availability

The training files and trained classifier used in this performance assessment, essential scripts, and input files for training the q2-feature-classifier plugin for use in QIIME2 are available at https://github.com/HillLabSask.

## References

1. Links MG, Dumonceaux TJ, Hemmingsen SM, Hill JE. The chaperonin-60 universal target is a barcode for bacteria that enables de novo assembly of metagenomic sequence data. PloS One. 2012;7: e49755. doi:10.1371/journal.pone.0049755

2. Links MG, Demeke T, Gräfenhan T, Hill JE, Hemmingsen SM, Dumonceaux TJ. Simultaneous profiling of seed-associated bacteria and fungi reveals antagonistic interactions between microorganisms within a shared epiphytic microbiome on Triticum and Brassica seeds. New Phytol. 2014;202: 542–553.

3. Verbeke TJ, Sparling R, Hill JE, Links MG, Levin D, Dumonceaux TJ. Predicting relatedness of bacterial genomes using the chaperonin-60 universal target (cpn60 UT): application to Thermoanaerobacter species. Syst Appl Microbiol. 2011;34: 171–179.

4. Shukla I, Hill JE. cpn60 barcode sequences accurately identify newly defined genera within the Lactobacillaceae. Can J Microbiol. 2022;68: 457–464. doi:10.1139/cjm-2021-0296

5. Schellenberg JJ, Paramel Jayaprakash T, Withana Gamage N, Patterson MH, Vaneechoutte M, Hill JE. Gardnerella vaginalis subgroups defined by cpn60 sequencing and sialidase activity in isolates from Canada, Belgium and Kenya. PLoS One. 2016;11: e0146510. doi:10.1371/journal.pone.0146510

6. Hill JE, Gottschalk M, Brousseau R, Harel J, Hemmingsen SM, Goh SH. Biochemical analysis, cpn60 and 16S rDNA sequence data indicate that Streptococcus suis serotypes 32 and 34, isolated from pigs, are Streptococcus orisratti. Vet Microbiol. 2005;107: 63–9.

7. Katyal I, Chaban B, Hill JE. Comparative genomics of cpn60 defined Enterococcus hirae ecotypes and relationship of gene content differences to competitive fitness. Microb Ecol. 2015;72: 917–930. doi:doi 10.1007/s00248-015-0708-2

8. Paramel Jayaprakash T, Schellenberg JJ, Hill JE. Resolution and characterization of distinct cpn60-based subgroups of Gardnerella vaginalis in the vaginal microbiota. PLoS ONE. 2012;7: e43009. doi:10.1371/journal.pone.0043009

9. Tian Q, Zhao W, Lu S, Zhu S, Li S. DNA Barcoding for efficient species- and pathovar-level identification of the quarantine plant pathogen Xanthomonas. PLoS One. 2016;11: e0165995. doi:10.1371/journal.pone.0165995

10. Hill JE, Town JR, Hemmingsen SM. Improved template representation in cpn60 PCR product libraries generated from complex templates by application of a specific mixture of PCR primers. Environ Microbiol. 2006;8: 741–746. doi:doi:10.1111

11. Goh SH, Potter S, Wood JO, Hemmingsen SM, Reynolds RP, Chow AW. HSP60 gene sequences as universal targets for microbial species identification: studies with coagulasenegative staphylococci. J Clin Microbiol. 1996;34: 818–23.

12. Vancuren SJ, Hill JE. Update on cpnDB: a reference database of chaperonin sequences. Database. 2019;2019: doi:10.1093/database/baz033. doi:10.1093/database/baz033

13. Chaban B, Musil KM, Himsworth CG, Hill JE. Development of cpn60-based real-time quantitative PCR assays for the detection of 14 Campylobacter species and application to screening canine fecal samples. Appl Environ Microbiol. 2009;75: 3055–3061.

14. Rohde J, Rubin JE, Kulathunga DGRS, Hill JE, Habighorst-Blome K, Hampson DJ, et al. Identification of Brachyspira species by cpn60 universal target sequencing is superior to NADH oxidase gene sequencing. Vet Microbiol. 2019;239: 108454. doi:10.1016/j.vetmic.2019.108454

15. Comte A, Grafenhan T, Links MG, Hemmingsen SM, Dumonceaux TJ. Quantitative molecular diagnostic assays of grain washes for Claviceps purpurea are correlated with visual determinations of ergot contamination. PLoS One. 2017;12: e0173495. doi:10.1371/journal.pone.0173495

16. Bai L, Paek J, Shin Y, Park H-Y, Chang YH. Lentilactobacillus kribbianus sp. nov., isolated from the small intestine of a mini pig. Int J Syst Evol Microbiol. 2020;70: 6476–6481. doi:10.1099/ijsem.0.004560

17. Joshi A, Thite S, Karodi P, Joseph N, Lodha T. Alkalihalobacterium elongatum gen. nov. sp. nov.: An Antibiotic-Producing Bacterium Isolated From Lonar Lake and Reclassification of the Genus Alkalihalobacillus Into Seven Novel Genera. Front Microbiol. 2021;12. Available: https://www.frontiersin.org/article/10.3389/fmicb.2021.722369

18. Sakamoto M, Ikeyama N, Kunihiro T, Iino T, Yuki M, Ohkuma M. Mesosutterella multiformis gen. nov., sp. nov., a member of the family Sutterellaceae and Sutterella megalosphaeroides sp. nov., isolated from human faeces. Int J Syst Evol Microbiol. 2018;68: 3942–3950. doi:10.1099/ijsem.0.003096

19. Chaban B, Albert A, Links MG, Gardy J, Tang P, Hill JE. Characterization of the upper respiratory tract microbiomes of patients with pandemic H1N1 influenza. PLoS ONE. 2013;8: e69559. doi:10.1371/journal.pone.0069559

20. Elwood C, Albert AYK, McClymont E, Wagner E, Mahal D, Devakandan K, et al. Different and diverse anaerobic microbiota were seen in women living with HIV with unsuppressed HIV viral load and in women with recurrent bacterial vaginosis: a cohort study. BJOG Int J Obstet Gynaecol. 2019;127: 250–259. doi:10.1111/1471-0528.15930

21. Dos Santos SJ, Pakzad Z, Elwood CN, Albert AYK, Gantt S, Manges AR, et al. Early Neonatal Meconium Does Not Have a Demonstrable Microbiota Determined through Use of Robust Negative Controls with cpn60-Based Microbiome Profiling. Microbiol Spectr. 2021;0: e00067–21. doi:10.1128/Spectrum.00067-21

22. Peterson SW, Knox NC, Golding GR, Tyler SD, Tyler AD, Mabon P, et al. A Study of the Infant Nasal Microbiome Development over the First Year of Life and in Relation to Their Primary Adult Caregivers Using cpn60 Universal Target (UT) as a Phylogenetic Marker. PloS One. 2016;11: e0152493. doi:10.1371/journal.pone.0152493

23. McKenney EA, Ashwell M, Lambert JE, Fellner V. Fecal microbial diversity and putative function in captive western lowland gorillas (Gorilla gorilla gorilla), common chimpanzees (Pan troglodytes), Hamadryas baboons (Papio hamadryas) and binturongs (Arctictis binturong). Integr Zool. 2014;9: 557–569. doi:10.1111/1749-4877.12112

24. Costa MO, Chaban B, Harding JCS, Hill JE. Characterization of the fecal microbiota of pigs before and after inoculation with “Brachyspira hampsonii.” PLoS ONE. 2014;9: e106399.

25. Dumonceaux TJ, Hill JE, Pelletier C, Paice MG, Van Kessel AG, Hemmingsen SM. Molecular characterization of microbial communities in Canadian pulp and paper activated sludge and quantification of a novel Thiothrix eikelboomii-like bulking filament. Can J Microbiol. 2006;52: 494–500.

26. Links MG, Dumonceaux TJ, McCarthy EL, Hemmingsen SM, Topp E, Town JR. CaptureSeq: Hybridization-Based Enrichment of cpn60 Gene Fragments Reveals the Community Structures of Synthetic and Natural Microbial Ecosystems. Microorganisms. 2021;9: 816. doi:10.3390/microorganisms9040816

27. Town JR, Links MG, Fonstad TA, Dumonceaux TJ. Molecular characterization of anaerobic digester microbial communities identifies microorganisms that correlate to reactor performance. Bioresour Technol. 2014;151: 249–57. doi:10.1016/j.biortech.2013.10.070

28. Schellenberg J, Links MG, Hill JE, Dumonceaux TJ, Peters GA, Tyler S, et al. Pyrosequencing of the Chaperonin-60 Universal Target as a Tool for Determining Microbial Community Composition. Appl Environ Microbiol. 2009;75: 2889–2898. doi:10.1128/AEM.01640-08

29. Wang Q, Garrity GM, Tiedje JM, Cole JR. Naive Bayesian classifier for rapid assignment of rRNA sequences into the new bacterial taxonomy. Appl Environ Microbiol. 2007;73: 5261–7. doi:AEM.00062-07 [pii] 10.1128/AEM.00062-07

30. Liu K-L, Porras-Alfaro A, Kuske CR, Eichorst SA, Xie G. Accurate, rapid taxonomic classification of fungal large-subunit rRNA genes. Appl Environ Microbiol. 2012;78: 1523–1533. doi:10.1128/AEM.06826-11

31. Porter TM, Hajibabaei M. Automated high throughput animal CO1 metabarcode classification. Sci Rep. 2018;8: 4226. doi:10.1038/s41598-018-22505-4

32. Bell KL, Loeffler VM, Brosi BJ. An rbcL reference library to aid in the identification of plant species mixtures by DNA metabarcoding. Appl Plant Sci. 2017;5: apps.1600110. doi:10.3732/apps.1600110

33. Vancuren SJ, Santos SJD, Hill JE, Maternal Microbiome Legacy Project Team. Evaluation of variant calling for cpn60 barcode sequence-based microbiome profiling. PLOS ONE. 2020;15: e0235682. doi:10.1371/journal.pone.0235682

34. Meng G, Li Y, Yang C, Liu S. MitoZ: a toolkit for animal mitochondrial genome assembly, annotation and visualization. Nucleic Acids Res. 2019;47: e63. doi:10.1093/nar/gkz173

35. Huerta-Cepas J, Serra F, Bork P. ETE 3: Reconstruction, Analysis, and Visualization of Phylogenomic Data. Mol Biol Evol. 2016;33: 1635–1638. doi:10.1093/molbev/msw046

36. Ren Q. Characterization of the salivary microbiome in COVID-19 infection and development of a cpn60 classifier. University of Saskatchewan. 2023.

37. McClymont E, Albert AY, Wang C, Dos Santos SJ, Coutlée F, Lee M, et al. Vaginal microbiota associated with oncogenic HPV in a cohort of HPV-vaccinated women living with HIV. Int J STD AIDS. 2022; https://doi.org/10.1177/09564624221109686. doi:10.1177/09564624221109686

38. Dos Santos SJ, Pakzad Z, Albert AYK, Elwood C, Grabowska K, Links MG, et al. Impact of vaginal microbiota and delivery mode on early-life infant stool microbiomes. Submitted.

39. Johnson LA, Chaban B, Harding JC, Hill JE. Optimizing a PCR protocol for cpn60-based microbiome profiling of samples variously contaminated with host genomic DNA. BMC Res Notes. 2015;8: 253. doi:10.1186/s13104-015-1170-4

40. Silva reference files. In: https://mothur.org [Internet]. [cited 17 Oct 2022]. Available: https://mothur.org

41. Vancuren SJ, Santos SJD, Hill JE, Team the MMLP. Evaluation of variant calling for cpn60 barcode sequence-based microbiome profiling. PLOS ONE. 2020;15: e0235682. doi:10.1371/journal.pone.0235682

42. Werner JJ, Koren O, Hugenholtz P, DeSantis TZ, Walters WA, Caporaso JG, et al. Impact of training sets on classification of high-throughput bacterial 16s rRNA gene surveys. ISME J. 2012;6: 94–103. doi:10.1038/ismej.2011.82

43. Vargas-Albores F, Ortiz-Suárez LE, Villalpando-Canchola E, Martínez-Porchas M. Size-variable zone in V3 region of 16S rRNA. RNA Biol. 2017;14: 1514–1521. doi:10.1080/15476286.2017.1317912

